# High-throughput electrochemical transformation expands mass spectrometric reference space for transformation products

**DOI:** 10.64898/2026.11.14.765618

**Authors:** Erik Niehaves, Manuel Groß, Giovanni A. Vitale, Steffen Heuckeroth, Thilo Hofmann, Uwe Karst, Daniel Petras, Valentin Göldner

**Author notes:** Correspondence and requests for materials should be addressed to: Valentin Göldner –, Daniel Petras –.

## Abstract

Structure annotation of unknown features detected by high-resolution tandem mass spectrometry remains challenging. Spectral libraries provide confident annotation but have limited chemical space coverage. This hampers discovery of natural products and xenobiotics and often necessitates laborious isolation and characterization. We introduce a scalable high-throughput workflow combining online electrochemistry with high-resolution tandem mass spectrometry to expand the empirical spectral reference space for transformation products. This approach enables generating thousands of potential transformation products used for prioritization in non-target datasets. Applying the workflow to 112 chemicals, we generated a suspect spectral library containing 1,341 transformation product entries. Repository-scale matching against public non-target datasets yielded matches for 214 electrochemical transformation products with 70% not annotated by conventional open-access spectral libraries. This workflow provides a source of empirical reference spectra for overlooked transformation products, extends coverage beyond compounds represented in conventional libraries, and supports prioritization of features for structural elucidation, quantification, and bioactivity assessment.

## 1. Introduction

Oxidation and reduction are central processes governing chemical transformations across natural and engineered environments. In humans, animals, microbes, and plants, redox reactions are central metabolic processes, increasing the polarity of natural products and xenobiotics or introducing functional groups for further modification.^1^ In engineered systems, redox processes drive the transformation and degradation of xenobiotics during wastewater treatment^2^ and the aging of synthetic materials like rubber.^3^ These transformations result in a broad chemical space of transformation products with diverse physico-chemical properties. For natural products, these processes expand chemical diversity and may give rise to derivatives with altered bioactivity,^4^ whereas transformation products of environmental contaminants may exhibit increased persistence, mobility, or toxicity compared to their precursor compounds.^5,6^ Nevertheless, the vast majority of transformation products remain inaccessible as authentic physical standards and inherently underrepresented in reference databases such as tandem mass spectral libraries. This results in an analytical bottleneck for both the discovery of new biologically active natural products and the holistic monitoring of potentially hazardous xenobiotics.^6^

Non-target analysis using liquid chromatography and high-resolution tandem mass spectrometry (LC-HRMS/MS) has proven to be a powerful tool for characterizing metabolic profiles and detecting xenobiotics at trace level. However, confident identification of features detected during non-target studies remains challenging due to limited availability of authentic standards and high-quality reference data.^7,8^ Chemical synthesis and isolation of transformation products from complex matrices are slow and laborious. Therefore, these efforts are focused only on a few prioritized features, limiting discovery at a broader scale.

Electrochemical simulation of redox transformations combined with online mass spectrometric characterization (EC-MS) offers an alternative route to generate empirical reference data without requiring authentic standards for each transformation product.^9^ EC-MS has been used since the late 1990s and early 2000s to simulate oxidative and reductive human phase I metabolism^10–12^ and environmental transformation processes.^13–15^ However, throughput has been limited by instrumental setups designed primarily for single-compound analysis and a lack of automated data processing, requiring laborious manual annotation of electrochemical transformation products (ETPs). Here, we introduce a scalable high-throughput workflow for the acquisition, annotation, and curation of empirical HRMS/MS reference data for ETPs. By integrating an automated post-column electrochemical reaction with an adapted data processing workflow using feature shape and retention time correlation^16^ in mzmine,^17,18^ we increased the throughput of ETP generation and reference data curation compared with conventional EC-MS workflows. We applied this approach to xenobiotic and natural product standards to generate a publicly accessible suspect spectral library. To assess the occurrence of the ETPs, we matched the product spectra against public non-target datasets at the repository scale totaling more than 1 billion MS/MS spectra from thousands of independently acquired HRMS/MS datasets.^19^ Together, our results highlight that electrochemical transformation can be scaled into a systematic source of empirical library spectra and thereby expand the reference space available for prioritizing novel transformation products in non-target mass spectrometry (Figure 1).

**Figure 1.**
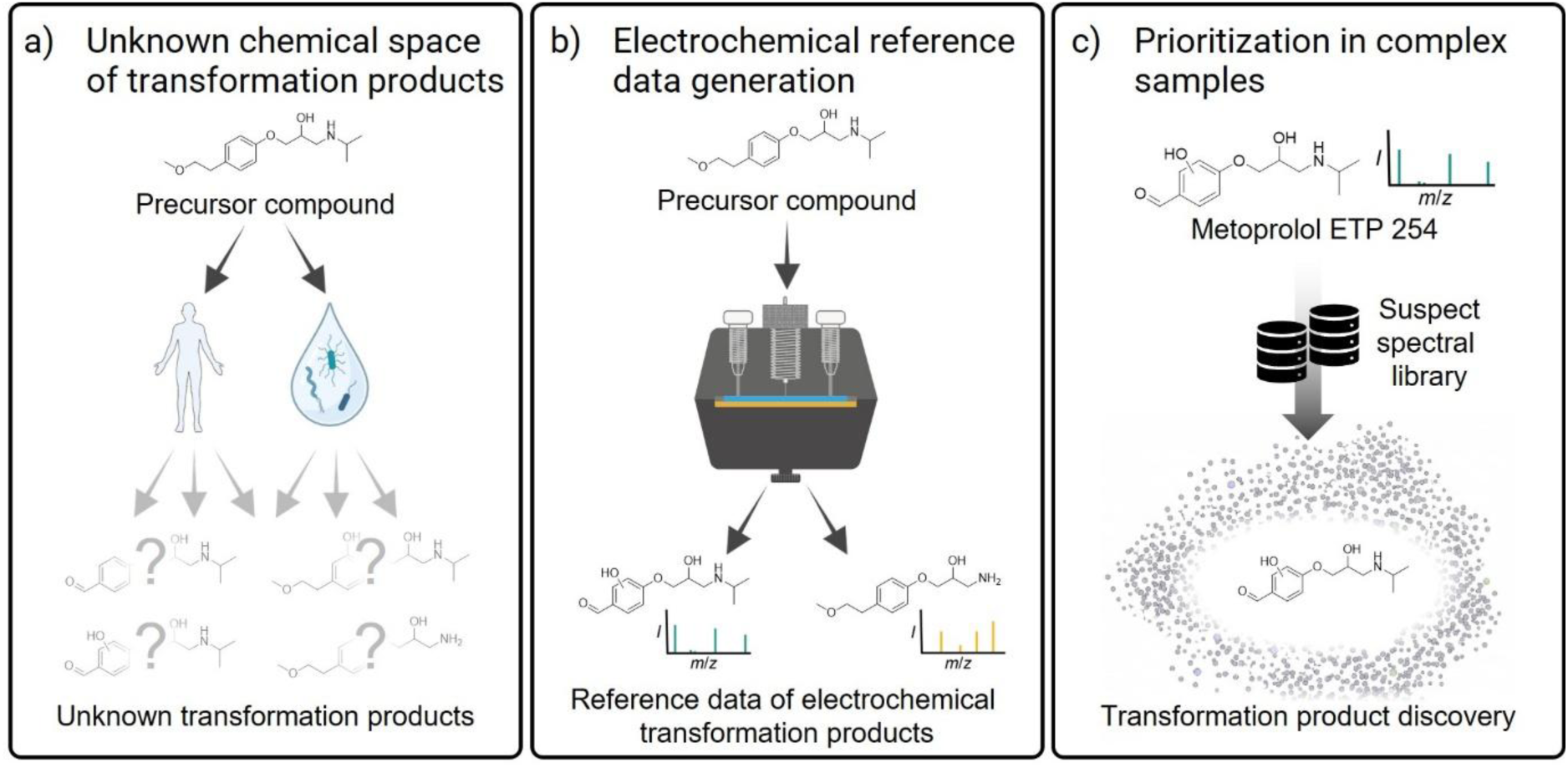
A scalable, automated electrochemistry workflow to explore the unknown chemical space of transformation products. a) Redox reactions in natural and engineered environments generate transformation products that are often uncharacterized, creating an analytical bottleneck. b) Automated electrochemical simulation systematically produces electrochemical transformation products (ETPs) and acquires empirical high-resolution tandem mass spectra at high throughput. c) The resulting suspect spectral library enables the prioritization of previously uncharacterized transformation products in complex non-target datasets. Parts of this figure were created with BioRender.com.

## 2. Results

### Scalable generation of empirical ETP spectra using LC-EC-HRMS/MS

Throughput of EC-MS data acquisition was increased by analyzing mixtures of precursor compounds in a single run. Online hyphenation of liquid chromatography with EC-MS enables the alignment of ETPs and precursor compounds based on retention time and peak shape.^20^

System suitability was evaluated by analyzing a set of eight compounds with known transformation behavior in online EC-MS.^12,15,21–29^ Conversion efficiency was investigated using the relative chromatographic peak area of the precursor compounds between untransformed analyses and electrochemical conversion at different cell potentials (Figure 2a and c). Assignment of ETPs to related precursor compounds based on chromatographic coelution was evaluated by extracting ion chromatograms of precursors and ETPs based on literature (Figure 2b). The formation of expected ETPs was observed in the high-throughput setup and their coelution was verified, enabling automated ETP annotation.

**Figure 2.**
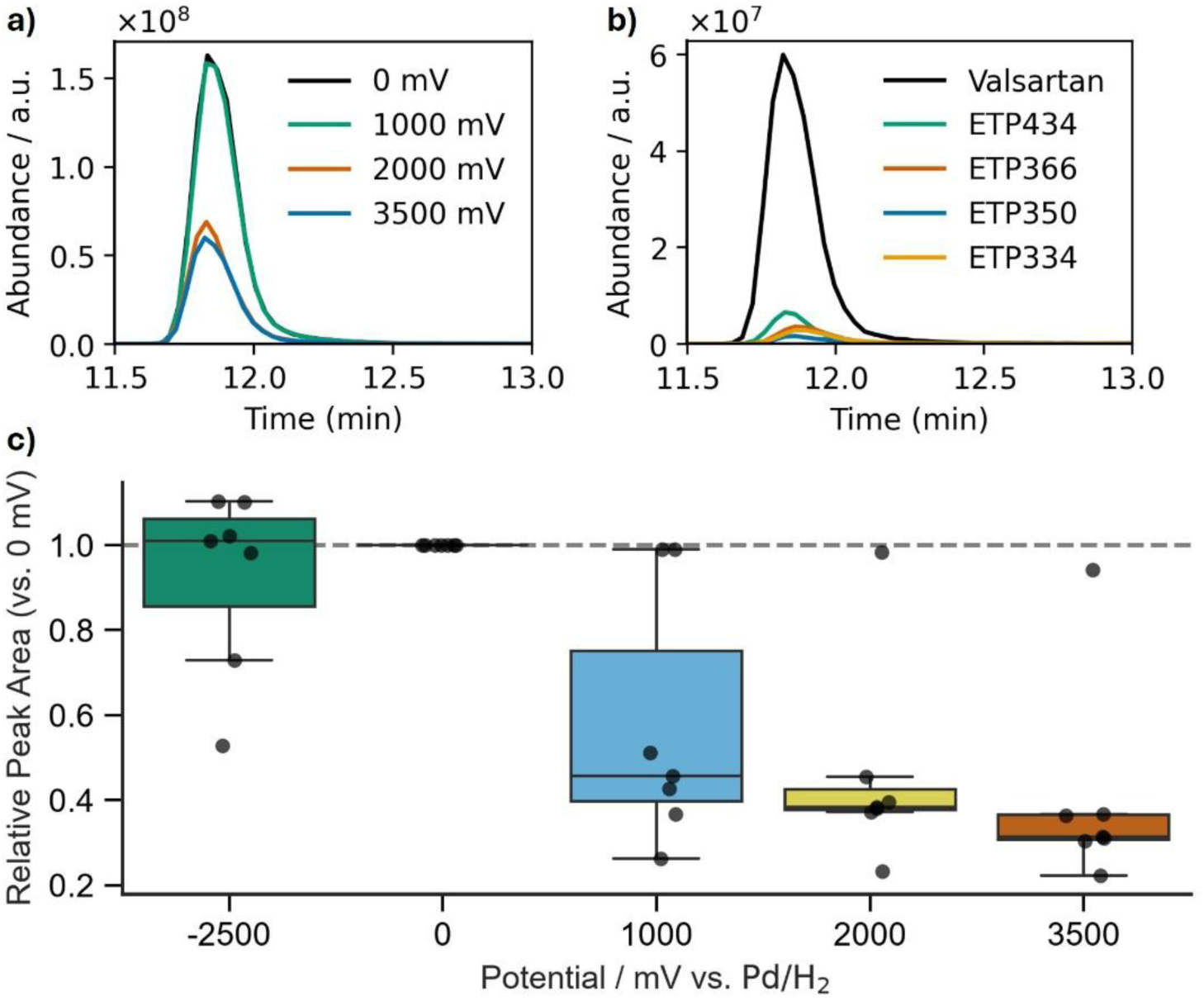
System suitability evaluation using model compounds with known electrochemical transformation behavior. a: Extracted ion chromatograms of Valsartan ([M+H]^+^, *m*/*z* 436.2343) at different electrochemical cell potentials using a boron-doped diamond (BDD) electrode. b: Extracted ion chromatograms of Valsartan and four electrochemical transformation products (ETPs) described in literature^29^ at 2000 mV on a BDD electrode. ETPs are named based on their nominal *m*/*z*. c: Peak areas of the seven precursor compounds detectable in positive ion mode at oxidative and reductive potentials relative to 0 mV.

Relative peak areas decrease with increasing cell potential down to a median relative peak area of 31% at 3500 mV across the seven compounds detected in positive ion mode. Only the sulfonylurea herbicide tritosulfuron shows minimal to no reduction of peak area at oxidative or reductive potential. This is in line with low yields of ETPs in literature even under optimized conditions.^28^

### Automated annotation of ETPs synchronizes data evaluation with high-throughput screening

High-throughput conversion acquires data at a rate where manual annotation of ETPs is not feasible. Thus, an automated data analysis workflow for post-column reactions was integrated in the open-source platform mzmine.^17^ Peak shape correlation was employed to group features with similar retention times and peak shapes.^16,18^ The resulting groups are the basis for ETP annotation using the “post-column reaction” processing module. The mzmine processing workflow is visualized in Figure 3.

**Figure 3.**
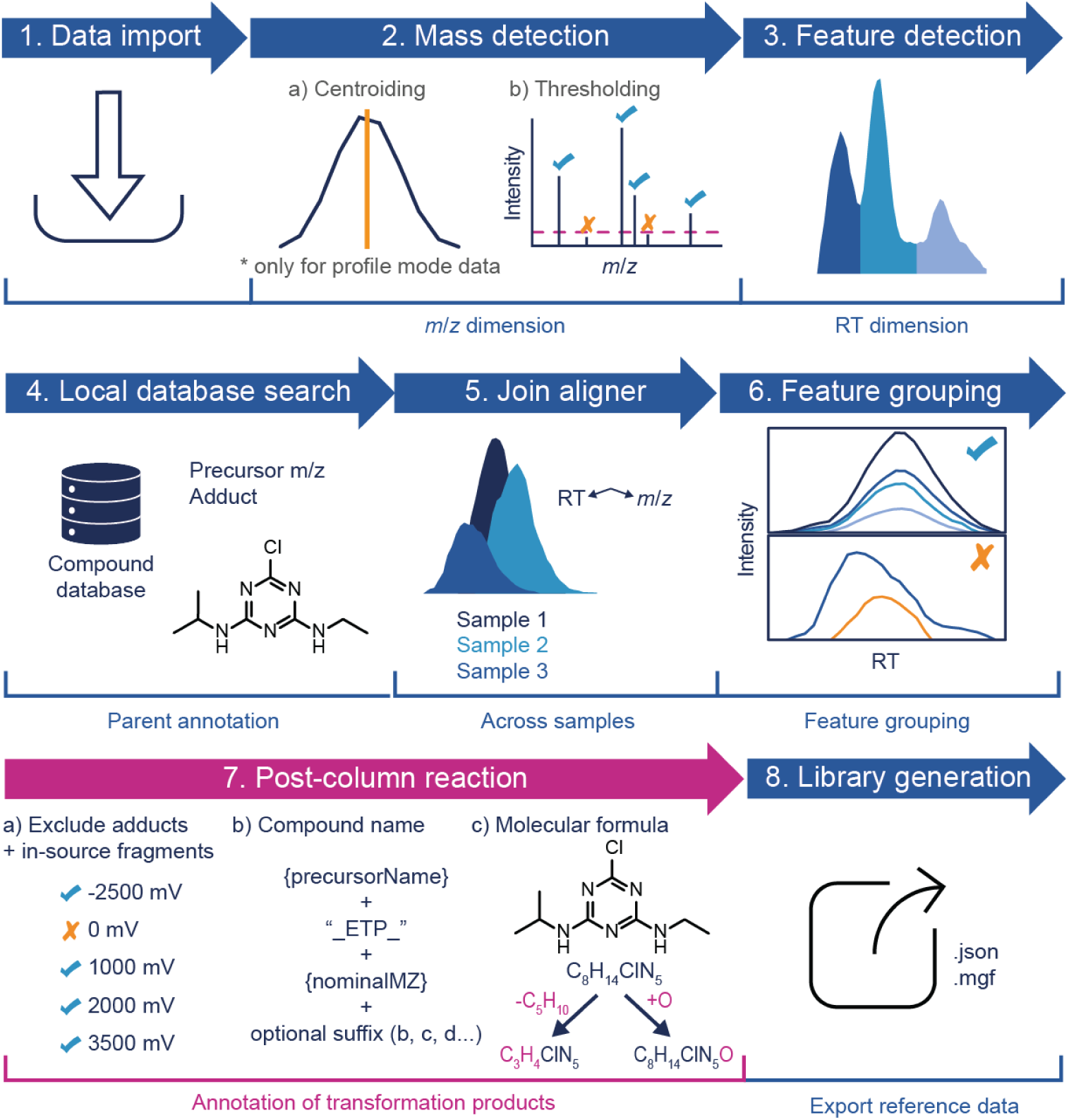
Processing workflow for automated annotation and suspect spectral library generation of electrochemical transformation products (ETPs) from liquid chromatography-electrochemistry-mass spectrometry. The “post-column reaction” module developed in this work (magenta) automatically assigns and annotates ETPs and is integrated in an mzmine feature detection and spectral library generation workflow. Figure adapted from mzmine documentation.^30^

The module considers features with peak-shape correlation to precursor compounds as their respective potential ETPs. The workflow actively removes common false positives by excluding adducts and in-source fragments of precursor compounds based on unreacted control samples. Furthermore, the false positive rate is reduced by considering only ETPs that are assigned to a molecular formula that is chemically plausible based on the respective precursor’s formula. These annotations are exported to a suspect spectral library^31^, if related HRMS/MS spectra meet defined quality criteria.

To benchmark the workflow, we first applied it to the system suitability compounds with well-documented transformations. The automated process successfully generated and annotated ETPs for 6 out of 7 oxidative reaction types expected from literature. None of the expected reductive transformations were observed under the applied conditions. Repeatability was assessed based on overlapping ETPs detected and annotated across repeated analyses within one day (80-82% repeatable) and across days (51-66% repeatable). Full validation results, including a detailed description of intra- and inter-day repeatability, are summarized in the supporting information (S1).

### High-throughput acquisition efficiently builds empirical reference data of redox transformation products

The high-throughput instrumental LC-EC-HRMS/MS setup was applied to screen the transformation of 112 precursor compounds (supporting information, S6). In addition to the previously discussed 8 compounds used for method optimization and evaluation, 87 xenobiotics (in mixtures containing 12 to 16 compounds each) and 17 natural products (in a single mix) were studied. Overall, 1,341 quality-filtered spectral entries assigned to suspect ETPs were generated and curated into suspect spectral libraries in approximately 37 hours of instrument time. This corresponds to an average of 12 ETPs per precursor compound and highlights the potential for high-throughput expansion of the chemical space of transformation products covered by spectral libraries. The resulting suspect spectral libraries for natural products and xenobiotics are accessible for download via Zenodo (https://doi.org/10.5281/zenodo.22960106).

### Reference data generated by electrochemical transformation provides spectral matches in public non-target data repositories

Public repositories of independently acquired non-target HRMS/MS data were used to approximate the detected chemical space available for retrospective analysis. We used this resource to assess whether spectral matches to the ETP library can be retrieved from public non-target datasets. This provides the first large-scale assessment of electrochemistry for the simulation of redox transformations across systems in a single study. Simultaneously, it provides a demonstration that precursor-linked ETP spectra retrieve signals not matched by the selected public libraries, indicating the value of a large-scale library of reference data for ETPs.

The ETP suspect spectral library was matched against MassIVE,^32^ Metabolomics Workbench,^33^ Metabolights,^34–36^ and the NORMAN Digital Sample Freezing Platform^37^ using the batch search functionality via the MASST workflow in GNPS2.^38^ Simultaneously, annotation with the GNPS spectral libraries^32,39^ indicates the novelty of information contained in the ETP suspect spectral library. An overview of the MASST search and open spectral library comparison is provided in Figure 4.

**Figure 4.**
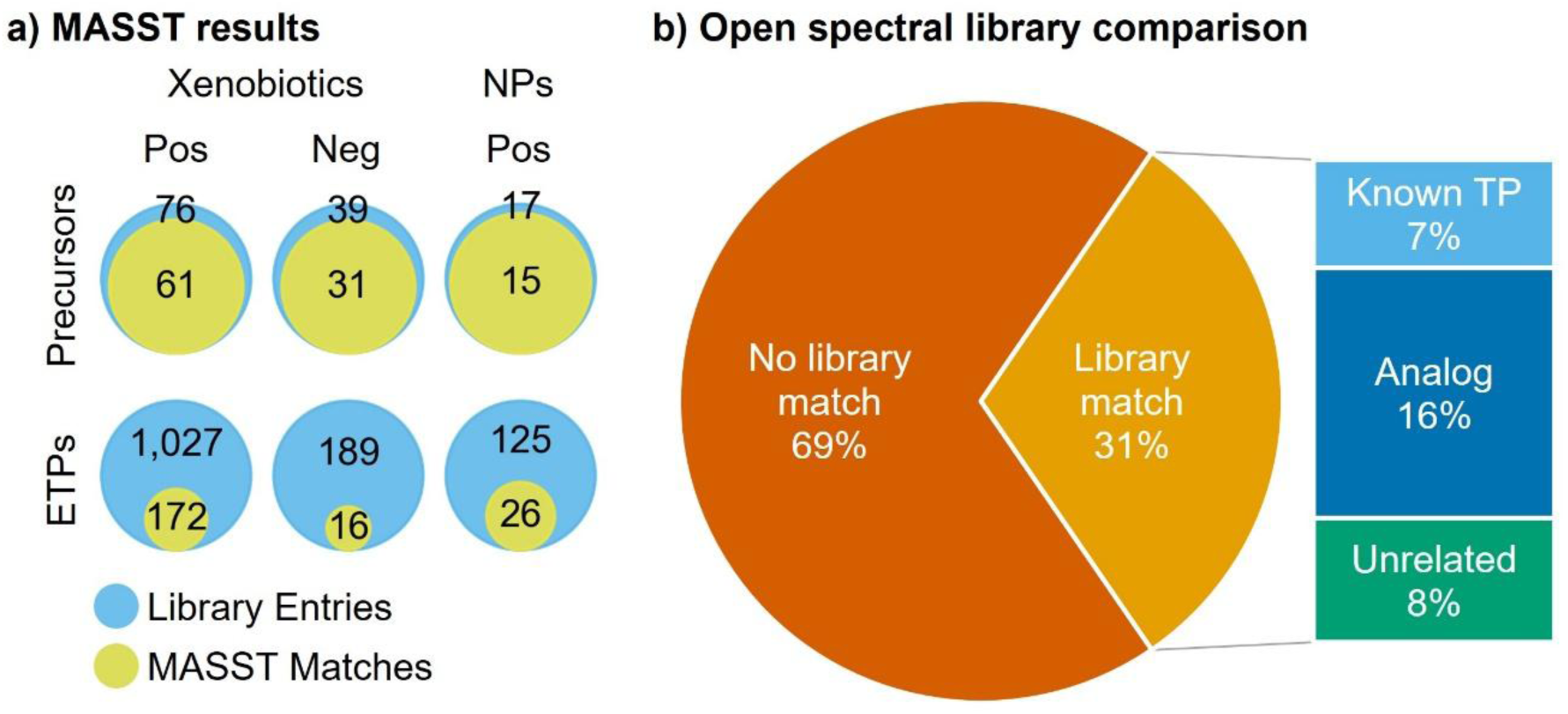
Overview of the MASST search and open spectral library comparison. a: MASST search results for xenobiotics and natural products displayed for precursor compounds and electrochemical transformation products (ETPs) separately. Blue: Total number of entries in the ETP suspect spectral library. Yellow: Number of entries that were matched in at least one open-access dataset. b: Open spectral library search results of the 188 xenobiotic ETPs matched during MASST search in positive and negative ion mode. Known TP: Matched with an open spectral library entry detailing a known transformation product of the ETP precursor compound; Analog: Matched with a structural analog of the ETP precursor compound; Unrelated: Structurally unrelated open spectral library match.

The reference library of xenobiotics and their ETPs contained data for 76 and 39 precursor compounds in positive and negative ion polarity, respectively. Out of these, 61 (80%, positive ion mode) and 31 (79%, negative ion mode) were matched to at least one spectrum in a public dataset across repositories. Reference data were acquired for 1027 ETPs in positive and 189 ETPs in negative ion mode. Across all repositories, 172 (17%) and 16 (8%) ETPs were annotated in at least one dataset in positive and negative ion mode, respectively (Figure 4a). 58 (31%) of these ETPs were additionally matched with spectra from the GNPS spectral libraries^32,39^ (Figure 4b). Entries matched both in public datasets and open spectral libraries were investigated for coherence of the respective spectral matches. 43 annotations explicitly refer to or are structurally similar to the ETP precursor compound across the public and ETP spectral libraries. Matches suggesting known transformation products support the use of electrochemistry to simulate biotic and abiotic transformation processes. In addition to spectral libraries containing exact structural information, the GNPS propagated suspect spectral library^39^ was employed. This library uses spectral network analysis to propose relations of unknown compounds to known molecules based on their spectral similarity. Common annotations of this propagated library and electrochemical simulation support each other by referring to the same precursor compound. Notably, 69% of the MASST matches of xenobiotic ETPs had no corresponding match in the tested open spectral libraries under the applied thresholds, indicating that they add new empirical reference data to the public domain.

For natural products, 15 out of 17 (88%) precursor compounds and 26 out of 125 (21%) ETPs were annotated in at least one open-access dataset. Among these, 7 ETPs were additionally annotated by the GNPS spectral libraries^32,39^ with 5 entries showing structural similarity to the ETP library precursor.

Exemplary ETPs were selected to discuss MASST and open library matches. Spectral mirror plots and proposed molecular structures are visualized in Figure 5.

**Figure 5.**
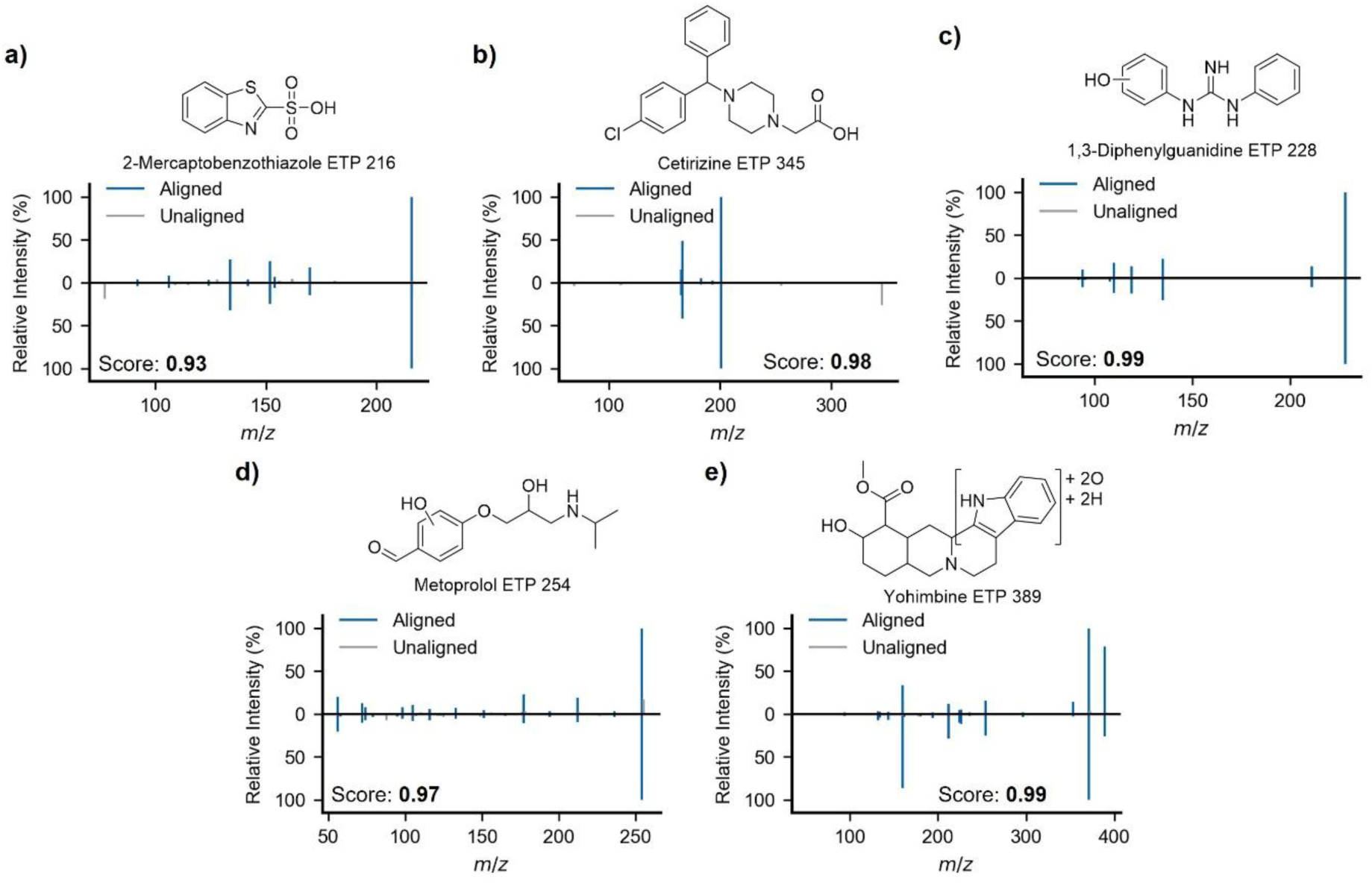
Exemplary matches of the ETP suspect spectral library in open-access datasets. All panels show the high-resolution tandem mass spectrum from the public dataset (top) and the corresponding ETP library spectrum (bottom), with proposed ETP structures. Proposed structures remain at candidate level and need to be verified by orthogonal characterization. a) 2-Mercaptobenzothiazole ETP 216 in dataset MSV000096860. b) Cetirizine ETP 345 in dataset MSV000092571. c) 1,3-Diphenylguanidine ETP 228 in dataset MSV000088021.^40^ d) Metoprolol ETP 254 in dataset MSV000091929.^41^ e) Yohimbine ETP 389 matched in MSV000096366.

All exemplary ETP matches show good spectral similarity and high cosine similarity scores above 0.90. Our suspect spectral library successfully confirmed known transformation products, as exemplified by 2-mercaptobenzothiazole ETP 216. This ETP spectrum matched the MassBank^42^ entry for its known sulfonic acid transformation product.^43^ Matches like this support the capability of electrochemical simulation to predict the formation of confirmed and extensively studied transformation products. Cetirizine ETP 345 was annotated by the propagated GNPS nearest neighbor suspect spectral library^39^ as “Related spectrum of ’Levocetirizine (Delta Mass:-44.026)” indicating relation to Levocetirizine, the R-enantiomer of Cetirizine. Both the propagated suspect library and the ETP library refer to the same precursor compound and therefore support each other in the annotation. This demonstrates the potential of electrochemical simulation to improve confidence of annotations derived from molecular networking by generating additional empirical evidence. 1,3-Diphenylguanidine ETP 228 is an example of a mismatch between electrochemical reference data and the propagated suspect spectral library^39^ which annotated this spectrum as “Conjugate of 1*H*-imidazo[4,5-c]pyridin-4-amine”. The ETP is annotated in 53 datasets, especially in environmental metabolomics datasets, e.g., of coral reef dissolved organic matter from Moorea, French Polynesia (MSV000088021^40^). In this case, the annotation by the ETP library is supported by higher spectral similarity and the presence of the precursor compound 1,3-diphenylguanidine in the same samples. Metoprolol ETP 254 was annotated exclusively by the ETP suspect spectral library. This ETP has a molecular formula consistent with a known carboxylic acid transformation product of metoprolol and bisoprolol, contained in suspect lists like SWISSPHARMA24.^44,45^ However, no HRMS/MS reference data of this transformation product are available in the tested spectral libraries and the fragment spectra annotated by the ETP suspect spectral library indicate the presence of an isomer formed by aromatic hydroxylation rather than carboxylic acid formation. Metoprolol ETP 254 was annotated in 40 public datasets with the highest matching score in a study of human urine metabolomics (MSV000091929), illustrating how the library can prioritize candidate drug-related features for follow-up. Finally, yohimbine ETP 389 was matched in 11 plant-derived datasets. The highest matching score was observed in leaf extract of *Rauvolfia vomitoria* (MSV000096366), which has been previously shown to produce yohimbine-like monoterpene indole alkaloids^46^ and a yohimbine spectral match was obtained in the same sample. While no unambiguous structure can be proposed based on current data, a likely site of the modification was proposed in Figure 5 using ModiFinder.^47^ ModiFinder results for this compound are shown in the supporting information (S5).

## 3. Discussion

Transformation products remain poorly represented in mass spectral libraries because authentic standards and experimental reference spectra are unavailable for many of these compounds limiting their annotation in non-target HRMS/MS data. This leaves potentially relevant transformation products unresolved in both newly acquired and existing datasets. Electrochemical transformation provides a route to generate empirical reference spectra from known precursor compounds, but conventional EC-MS approaches have lacked the throughput and automated data handling required for systematic library generation. Coupling high-throughput electrochemical transformation with automated spectral curation can address this gap across xenobiotic and natural product standards. The resulting suspect spectral libraries yielded matches across independently acquired public HRMS/MS datasets. Among the 58 xenobiotic ETP entries that matched both public datasets and open spectral libraries, 43 were associated with annotations referring either to known transformation products or to structures related to the respective precursor compound, providing an external consistency check for the electrochemically generated reference data. At the same time, 69% of the xenobiotic ETPs matched in public datasets had no corresponding match in the tested open spectral libraries. Our suggested workflow therefore adds empirical reference spectra that are not represented in these resources. This is particularly useful for retrospective analysis, because existing HRMS datasets can be searched without additional sample acquisition. The same library can also be used in future non-target studies to prioritize candidate transformation products for further investigation.

The natural product data extends this proof of concept beyond xenobiotics, with 26 natural product ETP library entries matching spectra in public metabolomics datasets. Due to the higher structural complexity of most natural products and the occurrence of isomers, HRMS/MS interpretation is less straightforward. Different isomers can share similar fragmentation patterns, and the present data therefore does not establish a specific metabolite annotation. Therefore, the main value of the ETP library in natural product discovery is not structural assignment, but the ability to link previously unresolved spectra to potentially structurally similar compounds. This forms the basis for discovery-driven hypothesis building and helps to select features for isolation and orthogonal characterization.

This distinction is central to how the library should be used. Electrochemically generated reference spectra are primarily a prioritization tool rather than a source of confident structural identification. Because transformation takes place after chromatographic separation, products formed from the same precursor reach the mass spectrometer without a second separation. While their shared elution profile is essential for linking ETPs to the precursor compound, it also means that isomeric or isobaric products may be co-isolated and contribute to chimeric HRMS/MS spectra. Finally, HRMS/MS data alone are often not sufficient to differentiate positional isomers. Thus, annotations remain candidate-level evidence and conclusions on compound identity, occurrence, exposure, or hazard require additional confirmation and, where relevant, quantitative and effect data.

The system-suitability experiments show where the present workflow performs well and where its coverage remains incomplete. 6 out of 7 expected oxidative reaction types were observed, whereas none of the three expected reductive reaction types was recovered under the general screening conditions. The present implementation is therefore better demonstrated for oxidative than for reductive transformations. Intra-day repeatability of quality-filtered ETP annotations was 80–82%, while the overlap between analyses performed on different days decreased to 51–66%. The reduction in inter-day overlap is attributed mainly to longer-term changes in instrument background and electrode surface chemistry affecting the relative abundance of ETPs and therefore their selection for fragmentation during data-dependent acquisition (DDA).

A useful feature of the approach is that prioritization is only the first step in a characterization pipeline. Prioritized candidates can be generated at larger scale by electrosynthesis to provide material for orthogonal characterization such as structure elucidation using nuclear magnetic resonance spectroscopy,^28^ for quantitative method development, and for biological testing. If isotopically labelled precursor compounds are available, the same principle may also provide labelled transformation product material for use as internal standards.^48^ Finally, the dataset of paired precursor and ETP spectra may be useful for computational approaches because even without complete structural assignments, they provide a structured set of precursor-product relationships from which spectral patterns can be learned.

Future work should focus on three areas: broader precursor-compound coverage, improved recovery of reductive and low-abundance products, and orthogonal validation of selected repository matches. Continued public release of these data would progressively enlarge the empirical reference space available for transformation products. The present workflow therefore provides a scalable experimental route from known precursor compounds to empirical ETP spectra that complement existing spectral libraries and help prioritize candidate transformation products occurring in independently acquired datasets for definitive chemical characterization.

## 4. Materials and Methods

### Chemicals

Acetonitrile (ACN, LC-MS grade), methanol (MeOH, LC-MS grade), and formic acid (FA, LC-MS grade) were obtained from Fisher Scientific. Dimethyl sulfoxide (DMSO, ≥ 99.9%) was purchased from Thermo Fisher Scientific. Nitric acid (65%; AnalaR NORMAPUR) was purchased from VWR International. Ultrapure water was produced using an ELGA PURELAB Chorus 1 system (ELGA LabWater, Veolia Water Technologies Deutschland). Individual stock solutions for xenobiotics were prepared from standards using appropriate solvents (i.e., ACN, MeOH, ultrapure water, DMSO, or ACN/ultrapure water). A list of all xenobiotic precursor compounds with information on molecular formula, SMILES, InChIKey, CAS number, vendor, purity, logP, and solvents used for preparation of stock solutions is provided in the supporting information (S6). LogP (computed by XlogP3 3.0^49^) was retrieved from PubChem (accessed: October 20, 2026).^50^ Xenobiotics were combined to form a system suitability mixture (i.e., QC mix) containing a set of eight compounds with literature-known electrochemical transformation behavior and six additional compound mixtures (mix 1-6). Compound combinations for mix 1-6 were selected based on maximum difference in logP values within groups to minimize coelution. All compounds were diluted for a final concentration of 5 µM in the mixtures using ultrapure water/ACN (90/10; *v*/*v*) as solvent. Solvent blanks consisted of ultrapure water/ACN (90/10; *v*/*v*).

Molecular formula, SMILES, InChIKey, CAS number, vendor, purity, and logP of natural products used in this study are provided in the supporting information (S6). All natural products were combined in one mixture.

### Liquid chromatography

Liquid chromatography for xenobiotics was performed on a Vanquish Horizon LC system (Thermo Fisher Scientific). Chromatographic separation was performed using a reversed-phase CORTECS Premier T3 VanGuard FIT column (2.1×100 mm, 2.7 μm particle size, 120 Å pore size, Waters) equipped with a CORTECS Premier T3 VanGuard FIT guard column cartridge (2.1 × 5 mm, 2.7 μm particle size, 120 Å pore size, Waters). Eluent A consisted of ultrapure water containing 0.1 vol.-% FA while ACN/ultrapure water (95/5; *v*/*v*) containing 0.1 vol.-% FA was used as eluent B. The flow rate was 100 μL/min, the column was kept at 40 °C and the injection volume was 5.0 μL. The applied gradient and further chromatographic parameters are described in the supporting information (S2).

Liquid chromatography for natural products was performed on a Vanquish Horizon LC system (Thermo Fisher Scientific). Chromatographic separation was performed using a reversed-phase Kinetex C18 column (2.1×150 mm, 1.7 μm particle size, 100 Å pore size, Phenomenex). Eluent A consisted of ultrapure water containing 0.1 vol.-% FA while ACN containing 0.1 vol.-% FA was used as eluent B. The flow rate was 100 μL/min, the column was kept at 25°C and the injection volume was 2.0 μL. The applied gradient and further chromatographic parameters are described in the supporting information (S2).

### Electrochemistry

A thin-layer electrochemical cell, µPrep-Cell 2.0 (Antec Scientific) was used for electrochemical transformation of both xenobiotics and natural products. The cell is based on a three-electrode setup, with a Pd/H_2_ pseudo reference electrode and a counter electrode made from conductive polyether ether ketone. All potentials are given relative to the Pd/H_2_ pseudo reference electrode. Xenobiotics were analyzed using working electrodes made of BDD and glassy carbon (GC) while natural products were analyzed exclusively on a BDD electrode. A custom-built potentiostat and software was used to operate the EC cell in potentiostatic mode. The BDD electrode was reactivated before sequences by cycling from −3000 mV to 3000 mV with 50 mV/s for one hour in 0.5 M nitric acid at a flow rate of 50 µL/min. The GC electrode was manually polished before analysis to remove potential depositions.

For online LC-EC-HRMS/MS, the effluent of the LC was introduced into the electrochemical cell for transformation before introduction to the HRMS (see supporting information (S2) for a schematic depiction). The mobile phase of the LC separation served as an electrolyte for electrochemical conversion. Hence, experimental conditions like flow rate, type and concentration of the supporting electrolyte salt, and concentration of the analytes were optimized to accommodate requirements for LC, EC, and MS. Conversion was screened by consecutive injections of the analytes at varying constant potentials. Discrete potentials were distributed across the accessible potential range determined by the type of working electrode. Xenobiotics were analyzed on BDD at 0 mV, 1000 mV, 2000 mV, 3500 mV, and −2500 mV and on GC at 0 mV, 1000 mV, 2500 mV, and −1500 mV. Natural products were analyzed on BDD at 0 mV, 1000 mV, 1500 mV, and 2000 mV. A detailed description of acquisition sequences is provided in the supporting information (S2).

### Mass spectrometry

Mass spectrometric detection was performed using an Orbitrap Exploris 240 high-resolution mass spectrometer (Thermo Fisher Scientific) for xenobiotic analyses while natural product analyses were conducted using a Q Exactive HF high-resolution mass spectrometer (Thermo Fisher Scientific). Both instruments were equipped with a heated electrospray ionization (HESI) source. Data for xenobiotics were acquired in alternating polarity while natural products were analyzed in positive ion mode. HRMS/MS data were acquired in DDA mode. For xenobiotics analyses, an Orbitrap resolution of 60,000 for HRMS scans and 22,500 for HRMS/MS scans was employed. The resolution for natural products analyses was 120,000 for HRMS and 15,000 for HRMS/MS scans. Higher-energy collision dissociation was used for fragmentation. Xenobiotics were analyzed at a defined cycle time of 1 s, an isolation window of *m*/*z* 0.8 and stepped normalized collision energy of 20%, 35%, 50%, and 65% with equal split times. Natural products DDA analyses were conducted as a Top5 experiment, an isolation window of *m*/*z* 1.0 and a stepped normalized collision energy of 25%, 35%, and 45% with equal split times. Detailed mass spectrometric acquisition parameters are provided in the supporting information (S2).

### Data processing

LC-EC-HRMS/MS data were converted into open mzML format using ThermoRawFileParserGUI (version 1.7.3).^51^ Settings for conversion are provided in the supporting information (S3). Feature detection and annotation of precursor compounds were performed using mzmine (version 4.9.27).^17^ Processing included data import, mass detection, chromatogram building,^52^ chromatogram smoothing,^53^ chromatogram blank subtraction, feature resolving, isotope filtering, local compound database search, join alignment, duplicate peak filtering, and correlation grouping.^16,18^ These steps were followed by the “post-column reaction” module developed in this work for the automated annotation of ETPs in LC-EC-HRMS/MS experiments. Finally, the annotated features with HRMS/MS spectra meeting defined quality criteria were exported to a suspect spectral library.^31^ Processing parameters were kept constant between xenobiotic and natural product analyses as well as between BDD and GC electrodes. Retention times of analytes and ETPs were less consistent between potentials when using a GC working electrode, which may be attributed to potential-dependent interaction of analytes with the electrode surface. Hence criteria for correlation grouping were stricter for BDD than for GC. Detailed parameters for each processing step and a description of the “post-column reaction” module are provided in the supporting information (S3 and S4) and in the mzmine processing batch file shared along with the mzML data files at (https://doi.org/10.5281/zenodo.22960737).

### Public repository and spectral library matching

Curated electrochemical reference data were exported to mgf format and matched against public repositories and spectral libraries using the batch search functionality of MASST.^38^ Queried repositories include MassIVE,^32^ Metabolomics Workbench,^33^ Metabolights,^34–36^ and the NORMAN Digital Sample Freezing Platform.^37^ Spectral matching was performed with the GNPS spectral libraries^32^ including the propagated nearest neighbor suspect spectral library.^39^ Matching parameters included minimum cosine similarity: 0.75; fragment *m*/*z* tolerance: 0.0010; precursor *m*/*z* tolerance: 0.0010; minimum matched signals: 4; analog search: disabled.

MASST search results were aggregated by merging results for ETP entries defined by molecular formula and name. The optional suffix in the name of ETPs used to differentiate isobars was neglected for merging because the suffix depends on the number of isobaric ETPs which may vary across electrodes. Instead, the molecular formula was used to differentiate isobars. Merging was performed for positive and negative ion mode independently, leaving the possibility for compounds to be detected in both modes. The three spectral library matches with the highest cosine similarity score were automatically extracted and added to the overview.

Coherence of open spectral library and ETP annotations was evaluated by manually comparing the top three spectral library matches with the ETP annotation and assigning a “Library Match Type”. Match type 1 relates to a spectral library match detailing a known transformation product of the ETP precursor compound. Type 2 indicates either a match with an entry in the propagated nearest neighbor suspect spectral library^39^ related to the ETP precursor compound or a spectral library entry with structural similarity to the ETP precursor compound. Finally, match type 3 is assigned to structurally unrelated annotations. Library match types are provided alongside the list of ETPs and MASST search results in the supporting information (S6).

## Data availability

All acquired LC-EC-HRMS/MS data files are available in mzML and raw format via Zenodo (https://doi.org/10.5281/zenodo.22960737) under the CC BY 4.0 license. The suspect ETP spectral libraries are available in mgf and json format as separate libraries for xenobiotics (ETPlib xeno) and natural products (ETPlib NP) via Zenodo (https://doi.org/10.5281/zenodo.22960106) under the CC BY 4.0 license.

## Code availability

The complete source code for the mzmine version containing the post-column reaction module is available at https://github.com/valentin-goeldner/mzmine via branch ‘PostColumnReaction’ under the MIT license. Documentation is available at https://github.com/valentin-goeldner/mzmine_documentation via branch ‘PostColumnReaction’ under the MIT license. The module will be implemented in the official release of mzmine. The complete source code and documentation for the latest mzmine version are available at https://github.com/mzmine/mzmine and https://github.com/mzmine/mzmine_documentation, respectively, under the MIT license. Aggregation of MASST search results was implemented in Python 3.10. Source code is available under the MIT license via https://github.com/valentin-goeldner/Aggregate-MASST-Batch-Results. Python scripts were generated using Gemini 2.5 pro and verified by the authors.

## Acknowledgements

The authors would like to thank Lea Ann Dailey and Gabriela Hädrich (both University of Vienna) for providing access to standards of selected xenobiotic precursor compounds.

## Author contributions

**E.N.**: Conceptualization, Methodology, Formal analysis, Investigation, Data curation, Writing – Review & Editing; **M.G.**: Methodology, Software, Validation, Formal analysis, Investigation, Data curation, Writing – Review & Editing, Visualization; **G.A.V.**: Conceptualization, Methodology, Investigation, Writing – Review & Editing; **S.H.**: Methodology, Software, Writing – Review & Editing; **T.H.**: Resources, Writing – Review & Editing, Supervision; **U.K.**: Conceptualization, Resources, Writing – Review & Editing, Supervision; **D.P.**: Conceptualization, Resources, Writing – Review & Editing, Supervision; **V.G.**: Conceptualization, Methodology, Software, Validation, Formal analysis, Investigation, Data curation, Writing – Original draft, Writing - Review & Editing, Visualization, Supervision, Project administration

## Supplementary Information

Supplementary Information is available for this paper.

