## Supporting information S1-S5 for "High-throughput electrochemical transformation expands mass spectrometric reference space for transformation products"

*Correspondence:

### S1: System suitability test using eight compounds with literature-known electrochemical transformation

A system suitability mix consisting of eight compounds with well-documented electrochemical transformation was used to benchmark the scalable liquid chromatography-electrochemistry-high-resolution tandem mass spectrometry (LC-EC-HRMS/MS) workflow from generation of electrochemical transformation products (ETPs) through data acquisition to annotation and curation of ETP suspect spectral libraries. Data were evaluated with respect to intra- and inter-day repeatability of annotated ETPs and comparison of observed ETPs to those reported in literature.

#### Quality criteria for ETPs used to determine repeatability

ETPs were considered for repeatability evaluation and literature comparison if they met defined quality criteria. First, only ETPs with a molecular formula predicted by the "post-column reaction" annotation workflow (see S4) were considered. Furthermore, at least one HRMS/MS spectrum needed to be acquired for an ETP to be considered. For inter-day repeatability and literature comparison, HRMS/MS spectra additionally needed to pass spectral quality criteria. These criteria were chosen to be consistent with the filters applied for ETP library generation.

#### Intra-day repeatability

Intra-day repeatability was determined by replicate analysis of the system suitability mix in ten consecutive analyses within one analysis sequence. An ETP was considered to be formed and annotated repeatably if it fulfilled the quality criteria in at least five out of ten replicates. Repeatability was determined for boron-doped diamond (BDD) and glassy carbon (GC) electrodes separately and yielded 80% and 82% repeatable ETPs for BDD and GC, respectively (see table S1.1). Occurrence of about 20% unrepeatable ETPs is primarily attributed to the requirement for HRMS/MS spectra. Reproducible fragmentation of the same precursor ions across analyses in data-dependent acquisition (DDA) is complicated by many ETPs co-eluting at similar and low abundance.

#### Inter-day repeatability

Inter-day repeatability was determined by comparing the results of the ten replicates used for intra-day repeatability to triplicates of the system suitability mix analyzed throughout the sequence of 87 other xenobiotics. Features meeting all quality criteria, including HRMS/MS spectral quality filters showed a 51% and 66% overlap for BDD and GC analyses, respectively (see table S1.1). The reduced overlap compared to intra-day repeatability is primarily attributed to low abundant, co-eluting ETPs being picked irreproducibly for fragmentation by DDA. The more pronounced reduction in repeatability on the BDD electrode is consistent with the higher number of ETPs it generates, which increases competition for DDA fragmentation. Additionally, long-term drift contributes to reduced inter-day comparability. This includes changes in instrument background noise and electrode surface chemistry.

Overall, repeatability is sufficient for screening ETPs in an automated and scalable fashion. Repeatability tests indicate the potential to increase coverage of HRMS/MS information for low abundant ETPs by analysis of replicates across instrument days. This needs to be weighed against increased instrument time associated with replicate analysis.

**Table S1.1. Summary of ETP annotation repeatability on boron-doped diamond (BDD) and glassy carbon (GC) electrodes.** Intra-day repeatability is the percentage of quality-filtered ETPs detected in a minimum of 5 out of 10 replicate analyses. Inter-day repeatability is the percentage of quality-filtered ETPs shared between analyses conducted on different days.

| | BDD | GC |
|---|---|---|
| Intra-day repeatability | 80% | 82% |
| Inter-day repeatability | 51% | 66% |

**Literature comparison**

The ETPs generated from the system suitability mix were compared to a curated list of transformations reported in literature.[1–11] This comparison was performed at two levels: the simulation of broad reaction classes and the detection of specific, individual ETPs.

First, the workflow's ability to simulate expected oxidative reaction types was assessed. The automated process successfully simulated and generated reference data for six out of seven expected overall oxidative reaction types (Table S1.2). The only oxidative reaction type not observed was the oxidation of an aldehyde to a carboxylic acid, a transformation known to have low electrochemical yields but described in literature to occur for Metoprolol.[1] In contrast, none of the three expected reductive reaction types were observed under these general screening conditions.

This is attributed to the low inherent reactivity of the reducible precursor compounds (e.g., tritosulfuron[10]) and the use of a non-specialized workflow, which differs from the optimized, single-compound conditions used in targeted literature studies. This result validates the platform's primary strength as a high-throughput tool for simulating oxidative transformation while reductive transformation requires further investigation and optimization.

Second, at a more granular level, we compared the specific molecular formulae of all detected ETPs against those in the literature database. Overall, the workflow detected 49% of all ETPs reported across both oxidative and reductive literature studies. The lower detection rate for individual products compared to reaction classes reflects the different experimental priorities between this general-purpose screening method and literature studies optimized for single precursor compounds.

**Table S1.2. Comparison of literature-expected electrochemical transformation types and transformations observed in this study.**

| Expected reaction | Type | Observed |
|---|---|---|
| Hydroxylation | Oxidation | Yes |
| Dehydrogenation | Oxidation | Yes |
| *O*-dealkylation | Oxidation | Yes |
| *N*-dealkylation | Oxidation | Yes |
| *S*-oxidation | Oxidation | Yes |
| Oxidative C-C cleavage | Oxidation | Yes |
| Aldehyde oxidation | Oxidation | No |
| Nitro reduction | Reduction | No |
| Hydrodefluorination | Reduction | No |
| Hydrogenation | Reduction | No |

### S2: Parameters for LC-EC-HRMS/MS data acquisition

#### Liquid chromatography

Chromatographic separation for xenobiotic analyses was performed on a Vanquish Horizon HPLC system equipped with a VH-A10-A02 autosampler, a VH-P10-A-02 pump module, and a VH-C10-A03 column compartment (all from Thermo Fisher Scientific). A CORTECS Premier T3 VanGuard FIT column (2.1×100 mm, 2.7 µm particle size, 120 Å pore size, Waters) equipped with an CORTECS Premier T3 VanGuard FIT guard column cartridge (2.1 × 5 mm, 2.7 µm particle size, 120 Å pore size, Waters) was used. Eluent A consisted of ultrapure water containing 0.1 vol.-% formic acid (FA) while acetonitrile (ACN)/ultrapure water (95/5; *v*/*v*) containing 0.1 vol.-% FA was used as eluent B. The column was kept at 40 °C under forced air conditions and the injection volume was 5.00 µL. The next injection was prepared during the equilibration phase, 0.5 min before injection. The gradient is described in table S2.1. The effluent of the electrochemical cell was diverted to waste during the equilibration stage from 16.5 min to 0.1 min of the next analysis.

**Table S2.1. Chromatographic gradient for LC-EC-HRMS/MS analysis of xenobiotic precursor compounds.**

| *t* / min | −2.5 | −0.8 | −0.8 | 0.0 | 11.0 | 16.5 |
|---|---|---|---|---|---|---|
| Eluent B / % | 10 | 10 | 10 | 10 | 90 | 90 |
| Flow / µL/min | 400 | 400 | 100 | 100 | 100 | 100 |

Chromatographic separation for natural product analyses was performed on a Vanquish Horizon HPLC system (Thermo Fisher Scientific). A Kinetex C18 column (2.1×150 mm, 1.7 µm particle size, 100 Å pore size, Phenomenex) was used. Eluent A consisted of ultrapure water containing 0.1 vol.-% FA while ACN containing 0.1 vol.-% FA was used as eluent B. The column was kept at 25°C and the injection volume was 2.0 µL. The flow rate was set to 100 µL/min using the gradient described in table S2.2.

**Table S2.2. Chromatographic gradient for LC-EC-HRMS/MS analysis of natural product precursor compounds.**

| $t$ / min | 0.0 | 10.0 | 13.0 | 13.1 | 18.0 |
|---|---|---|---|---|---|
| Eluent B / % | 5 | 99 | 99 | 5 | 5 |

**Mass spectrometry**

Mass spectrometric detection for xenobiotic analyses was performed using an Orbitrap Exploris 240 high-resolution mass spectrometer (Thermo Fisher Scientific) equipped with a heated electrospray ionization (HESI) source. Alternating ion polarity was used to acquire data in positive and negative mode.

HESI source parameters were as follows: spray voltage (pos): 0.0 – 0.1 min: 0 V, 0.1 – 16.5 min: 3400 V; spray voltage (neg): 0.0 – 0.1 min: 0 V, 0.1 – 16.5 min: −2300 V; sheath gas: 25 arbitrary units; aux gas: 5 arbitrary units; sweep gas: 0 arbitrary units; ion transfer tube temperature: 320°C; vaporizer temperature: 70°C.

Orbitrap scan parameters were as follows: Orbitrap resolution (at *m*/*z* 200): 60,000; scan range: *m*/*z* 70 – 1000; RF Lens: 70 %; AGC target mode: Standard; normalized AGC target: 100% (1.0E6); maximum injection time mode: auto (calculated based on transient length); microscans: 1; data type: profile; lock mass correction: positive: *m*/*z* 214.08963, negative: *m*/*z* 226.97845, *m*/*z* 112.98559. The default charge state was 1. Mild trapping was enabled.

Data-dependent acquisition (DDA) was performed with a defined cycle time of 1 s using the following parameters: quadrupole isolation window: *m*/*z* 0.8; fragmentation type: higher-energy collision dissociation (HCD); normalized HCD energy: stepped (20%, 35%, 50%, and 65% with equal split times); Orbitrap resolution (at *m*/*z* 200): 22,500; scan range: auto; AGC target mode: custom; normalized AGC target: 120% (1.2E5); maximum injection time mode: custom (40 ms); microscans: 1; data type: centroid; lock mass correction: disabled.

Dynamic exclusion was set to 6 s after acquiring 1 HRMS/MS scan with a mass tolerance of 5 ppm (high) and 3 ppm (low) and isotope exclusion activated. A targeted exclusion mass list was generated by injection of an instrument blank before each sequence. Any *m*/*z* detected at a minimum absolute intensity of 5.0E5 and a minimum of 6 consecutive scans at a minimum intensity of 1.0E4 was added to an unscheduled exclusion list. Targeted mass exclusion was performed without intensity threshold and a mass tolerance of 3 ppm (high) and 2 ppm (low). Isotope exclusion was enabled and an intensity threshold for acquiring MS2 scans was set to 5.0E5.

Mass spectrometric detection for natural product analyses was performed using a QExactive HF high-resolution mass spectrometer (Thermo Fisher Scientific) equipped with a HESI source. Data were acquired in positive ion mode using DDA.

HESI source parameters were as follows: spray voltage (pos): 3500 V; sheath gas: 30 arbitrary units; aux gas: 10 arbitrary units; sweep gas: 1 L/min; ion transfer tube temperature: 250°C; probe heater temperature: 200°C.

Orbitrap scan parameters were as follows: Orbitrap resolution (at *m*/*z* 200): 120,000; scan range: *m*/*z* 150 – 1500; S-Lens RF level: 50; AGC target: 1.0E6; maximum injection time: 100 ms; microscans: 1; data type: profile.

DDA was performed for the five most intense precursor ions: quadrupole isolation window: *m*/*z* 1.0; fragmentation type: HCD; normalized HCD energy: stepped (25%, 35%, and 45% with equal split times); Orbitrap resolution (at *m*/*z* 200): 15,000; AGC target: 5.0E5; maximum injection time: 50 ms; microscans: 1; data type: profile.

**Instrumental setup for LC-EC-HRMS/MS**

The instrumental setup used for acquisition of LC-EC-HRMS/MS data is depicted schematically in Figure S2.1.

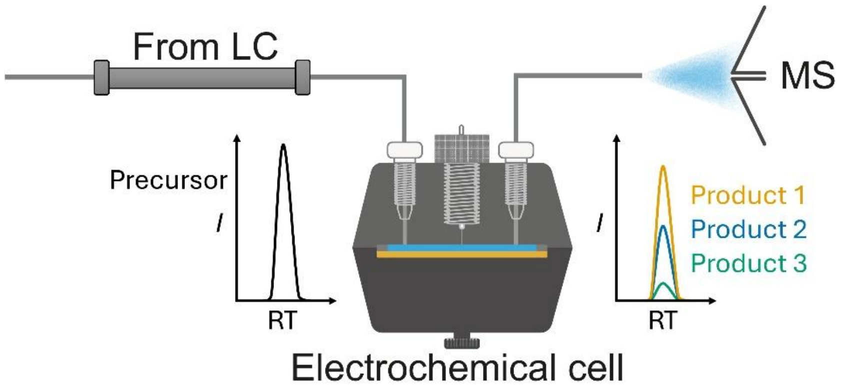


**Figure S2.1. Schematic depiction of the LC-EC-HRMS/MS setup.** The effluent of the LC is introduced into the electrochemical cell where conversion takes place. Transfer to the MS enables online detection and electrochemical transformation products can be assigned to their respective precursors by retention time and peak shape correlation.

### Acquisition sequence

LC-EC-HRMS/MS acquisition was performed by repeated injection of compound mixtures while applying selected constant electrochemical cell potentials per injection. Each compound mixture was analyzed by the same potential sequence before continuing to the next mixture. For xenobiotics on BDD, the potential sequence was: 0 mV, 1000 mV, 2000 mV, and 3500 mV, −2500 mV. Natural products were analyzed on BDD using the potential sequence: 0 mV, 1000 mV, 1500 mV, and 2000 mV. For xenobiotics analyses using GC electrodes, the potential sequence was: 0 mV, 1000 mV, and 2500 mV, −1500 mV.

Instrument banks consisting of ACN/ultrapure water (10/90, *v*/*v*) were analyzed for background subtraction. In the beginning of the acquisition, an instrument blank was analyzed using a complete potential sequence, followed by full potential sequences for the compound mixtures. A single instrument blank analysis at 0 mV was conducted between potential sequences for compound mixtures.

Because the analysis of xenobiotic ETPs comprised several compound mixtures, the performance of the acquisition system was monitored by repeated analyses of the compound mixture used for system suitability testing as quality control mixture (QC mix). The QC mix was analyzed using complete potential sequences in the beginning, middle, and end of the acquisition.

**S3: Parameters for LC-EC-HRMS/MS data processing**

HRMS/MS raw data were centroided and converted to mzML format using ThermoRawFileParserGUI (version 1.7.4).[12] The parameters for conversion were as follows: Spectrum (output file format): mzML; MS levels: all; peak picking: all; metadata: none; all detectors: no; exceptions: include reference and exception data; GZipped: no; compression: zlib; Errors: ignore missing instrument properties.

Mass detection was performed for MS1 and MS2 separately. The factor of lowest signal was used for mass detection using a noise factor of 5.0 (MS1) or 2.5 (MS2). Fragment scans were denormalized. FTMS shoulder peaks were filtered using the Lorentzian peak model function and a mass resolution of 60,000. After this step, positive and negative data were processed separately. Parameters differing for positive and negative modes are explicitly stated. For the chromatogram builder[13] the following parameters were set: all raw data files; scan filter: MS1, polarity: + (pos)/− (neg); minimum consecutive scans: 4; minimum intensity for consecutive scans: 1.0E4; minimum absolute height: 1.0E5; *m*/*z* tolerance (scan-to-scan): *m*/*z* 0.0020 or 10 ppm. Smoothing was applied to the feature lists generated in the previous step using the Savitzky Golay algorithm[14] with a retention time width of 11 scans. Chromatogram blank subtraction was applied, using all blanks and an *m*/*z* tolerance of *m*/*z* 0.0010 or 2.0 ppm. The local minimum feature resolver was applied to the feature lists generated in the previous step, using retention time as the resolving dimension. Parameters included: chromatographic threshold: 95.0%; minimum search range RT: 0.060 min; minimum relative height: 0.0%; minimum absolute height: 1.0E5; minimum ratio of peak top/edge: 1.50; peak duration range: 0.00-2.00 min; minimum scans: 4. The 13C isotope filter was used with an *m*/*z* tolerance of *m*/*z* 0.0010 or 2.0 ppm and a retention time tolerance of 0.04 min. Monotonic shape was required and the maximum charge was set to 2. The most intense isotope was used as the representative isotope. The isotopic peak finder used an *m*/*z* tolerance of *m*/*z* 0.0010 or 2.0 ppm and a maximum charge of isotope *m*/*z* of 1 searching in the single most intense scan. Local compound database search was performed before alignment to annotate precursor compounds only in samples where they were added. Matching was based on the precursor compound’s molecular formulae, considering $[M+H]^+$ and $[M-H]^-$ as adducts in positive and negative mode, respectively. The *m*/*z* tolerance was set to *m*/*z* 0.0005 or 2.0 ppm and the isotope matcher was enabled with an isotope *m*/*z* tolerance of *m*/*z* 0.0010 or 5.0 ppm, a minimum isotope intensity of 5% and a minimum isotope score of 0.7. The filename header was filtered for the column "unique_sample_id" which was used to annotate only precursor compounds spiked into the respective compound mixture. Feature alignment was performed using the join aligner on the feature lists generated by the previous step. Alignment parameters included: *m*/*z* tolerance: *m*/*z* 0.0005 or 2 ppm; weight for *m*/*z*: 3; retention time tolerance: 0.20 min; weight for RT: 1; Requirements for same charge state, same ID, isotope pattern similarity, and spectral similarity were disabled. Feature list rows were filtered allowing only features detected in a minimum of 1 sample, never removing features with MS2 scan or annotation. Duplicate peaks were filtered from the aligned feature list generated in the previous step using the "new average" filter mode using an *m*/*z* tolerance of *m*/*z* 0.0008 or 1.5 ppm, and a retention time tolerance of 0.20 min. Mobility tolerance and the requirement for identical identification was disabled. Correlation grouping was applied using the metaCorrelate module. Parameters differed between analyses made on BDD and GC electrodes. For BDD electrodes, an RT tolerance of 0.15 min, a minimum feature height of 1.0E5, and an intensity threshold for correlation of 1.0E4 were set. Minimum intensity overlap was set to 90% (xenobiotics) or 99% (natural products) and feature shape correlation was performed using cosine similarity algorithm with a minimum shape correlation of 40%, and a minimum of 5 overall data points and 2 data points on edge. For GC electrodes, an RT tolerance of 0.15 min, a minimum feature height of 1.0E5, and an intensity threshold for correlation of 1.0E4 were set. Minimum intensity overlap was set to 70%, and feature shape correlation was disabled.

The processing of the "post-column reaction" module is described in S4. The module was used to annotate ETPs using the feature list created by the previous step. Unreacted raw data files were selected based on the file name pattern containing the sub-string "_0mV". Molecular formula prediction was enabled with $[M+H]^+$ and $[M-H]^-$ as ionization types in positive and negative mode, respectively. The *m*/*z* tolerance for molecular formula prediction was *m*/*z* 0.0010 or 2.0 ppm and element heuristics rules H/C ratio, NOPS/C ratios and multiple element counts were enabled. Ring and double bond equivalents were restricted to −1 to 40 and only integer values. Isotope pattern filter and MS/MS filter were disabled. For BDD electrodes, a shape correlation threshold of 40% was applied while no shape correlation threshold was applied for analyses performed using the GC electrode. The name for automated transformation product annotation was set to "ETP".

Finally, reference data were exported to a spectral library in mzmine json format using batch spectral library generation.[15] Fragment scans were merged across samples with a tolerance of *m*/*z* 0.0020 or 5.0 ppm using the average intensity across scans. Chimeric spectra were determined by applying a minimum precursor purity of 60% using a target ion tolerance of *m*/*z* 0.0010 or 2.0 ppm and a precursor isolation tolerance of *m*/*z* 1.0. Chimeric spectra were included in the export and flagged. HRMS/MS quality criteria included a minimum number of 4 signals and a minimum explained intensity of 30%. Formula *m*/*z* tolerance was set to *m*/*z* 0.0010 or 5 ppm and only explained signals were exported.

### S4: Description of the “post-column reaction” module

To enable automated and scalable data processing for the LC-EC-HRMS/MS workflow, the “post-column reaction“ module was developed and integrated into the open-source platform mzmine.[16] This module is designed to systematically find and annotate transformation products (like ETPs) in post-column reaction experiments.

As ETPs are generated from a precursor compound after chromatographic separation, they are expected to co-elute, resulting in similar peak shapes and retention times. Additionally, ETPs should appear only in samples where a transformation was performed and be absent in unreacted control samples (i.e., analyses run at 0 mV). Therefore, a feature is considered a potential ETP if it meets both of these criteria: it must be grouped with a known precursor compound by peak correlation (metaCorrelate[17]), and it must be absent in all unreacted control runs.

The process is initiated with an aligned feature list containing data from both reacted and unreacted samples, in which precursor compounds have been identified through a prior database search or manual annotation. A prerequisite for the module is the pre-computation of chromatographic peak correlations across all features, using metaCorrelate. For each annotated precursor compound, the module queries its correlation group to identify features that are potential ETP candidates. These candidates are then systematically filtered. The first filter ensures that a candidate’s Pearson correlation score with its precursor meets an optional user-defined threshold (e.g., ≥40%). The second filter removes any candidate that is detected in one or more of the user-defined unreacted control samples. This eliminates background ions, adducts, and in-source fragments of the precursor compound, as these features would also be present in unreacted analyses.

Features that successfully pass these filtering steps are designated as ETP candidates and are automatically annotated. The compound name is systematically generated using the format {precursorName}_{tpName}_{nominalMZ}, where {precursorName} is inherited from the precursor, {tpName} is a String used for automated annotation of transformation products, and {nominalMZ} is the nominal *m*/*z* of the ETP. In this study, {tpName} was set to “ETP” To ensure unique identifiers for isobaric ETPs from the same precursor, an alphabetical suffix (e.g., _ETP_254b, _ETP_254c) is appended as needed.

To further increase annotation confidence and minimize false positives, the module incorporates an optional sub-task for molecular formula prediction. This step uses the known molecular formula of the precursor compound as a chemical constraint to narrow the search space for potential ETP formulae. The search space is automatically derived by allowing the elements contained in the precursor compound at a count from zero to the number of atoms of this element in the precursor. Common oxidative transformations that introduce additional atoms of selected elements, like hydroxylation and hydrogenation, are accounted for by increasing the upper limit for hydrogen and oxygen by eight. A candidate formula is assigned only if it matches the ETP's exact mass within a specified mass tolerance (e.g., ≤1 ppm) and meets additional optional criteria such as element count heuristics, ring and double bond equivalent restrictions, isotope pattern matching, and an MS/MS filter.

The final output of the module is the original feature list, modified in-place to include the newly generated ETP annotations and their predicted molecular formulae, providing a comprehensive and automatically curated dataset ready for subsequent analysis and spectral library generation.

**S5: Structural annotation and occurrence of yohimbine ETP 389**

The likely sites of modification to form yohimbine ETP 389 from yohimbine were determined by comparing HRMS/MS spectra from dataset MSV000096366 using ModiFInder.[18] Results for likely modification positions are depicted in Figure S5.1. Repository-scale search results are visualized in Figure S5.2 using the MASST search output.[19] Notably, yohimbine ETP 389 was matched exclusively in plant-derived samples with high ratios in genera *Raivolfia* and *Corynanthe*.

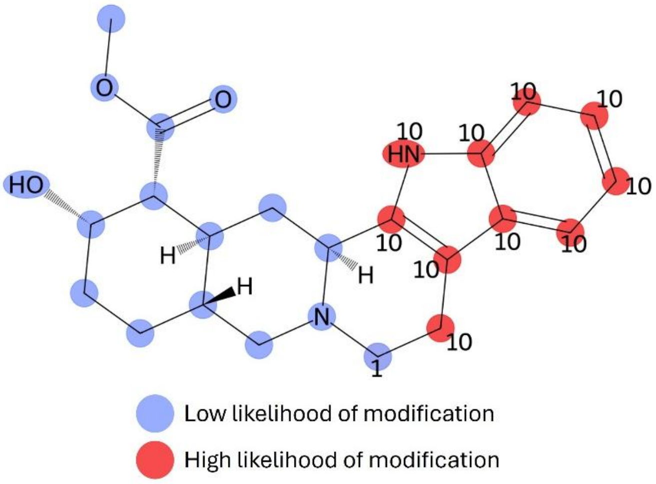


**Figure S5.1. ModiFinder[18] results for assigning modification locations between yohimbine and yohimbine ETP 389.** Assignment of likelihood of modification to form yohimbine ETP 389 from yohimbine. The likely sites of modification were determined using HRMS/MS spectra from dataset MSV000096366. Spectra used for comparison were: mzspec:MSV000096366:peak/pos/221116_LFX_OK_95_OK3_02_32_1_P1_ pos.mzML:scan:710 for yohimbine and mzspec:MSV000096366:peak/pos/221116 _LFX_OK_95_OK3_02_32_1_P1_pos.mzML:scan:416 for yohimbine ETP 389.

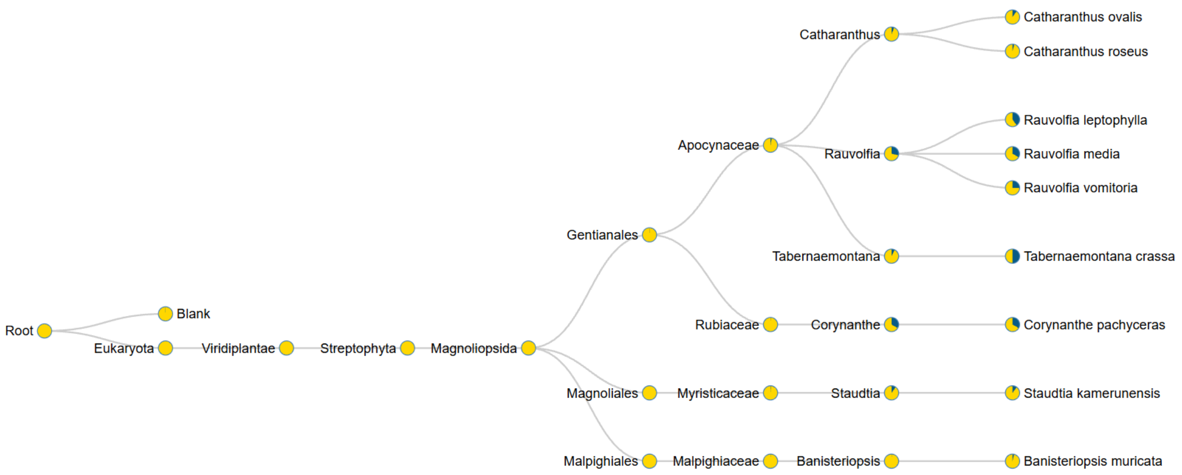


**Figure S5.2. Visualization of MASST[19] search results.** Pie charts display the proportion of samples with HRMS/MS spectral matches for yohimbine ETP 389 (blue fraction) compared to the total number of deposited samples of that type (yellow fraction).
